# Modulation of Ferroptosis During Early *Mycobacterium tuberculosis* Infection Contributes to Beijing Lineage Strain SA161 Virulence

**DOI:** 10.64898/2026.05.29.728786

**Authors:** Joshua J Ivie, Sylvia Stull, Ethan Bustad, Courtney R Plumlee, Sara B Cohen, Fergal J Duffy, Alan H Diercks, John D Aitchison, Kevin B Urdahl, Alexis Kaushansky, Alissa C Rothchild, Benjamin H Gern, Shuyi Ma

## Abstract

Although spread of internalized Mtb from the initial infected alveolar macrophages (AMs) is a crucial determinant of infection outcomes, the role of cell death in facilitating this spread, and how it is regulated by Mtb remains poorly understood. Ferroptosis, a lipid peroxidation-mediated form of necrotic cell death, contributes to pathology during later stages of infection. However, the role of ferroptosis in early AM cell death remains inadequately defined, and its induction during infection has been primarily studied with the laboratory strain, H37Rv. Using gene set variation analysis of single cell RNAseq data profiling Mtb-infected murine lungs, we found that the hypervirulent Beijing sublineage clinical strain SA161 is associated with an elevated pro-ferroptotic transcriptional response in AMs compared to H37Rv by 17 days post infection. Consistent with these transcriptional profiles, we found that SA161 induced increased lipid peroxidation in comparison to H37Rv during infection *in vitro* and *in vivo*. Administration of the lipid peroxidation inhibitor, ferrostatin-1 (Fer-1), reduces this Mtb-induced lipid peroxidation. Notably, we found that administration of Fer-1 to Mtb-infected mice significantly reduced bacterial burden for SA161 at 14 dpi while having no effect on H37Rv. Microscopic analysis of SA161-infected lung lesions at 14 dpi suggests that inhibition of ferroptosis-driving lipid peroxidation results in a greater proportion of AMs amongst infected cells and decreased neutrophil-associated IFN signaling. Collectively, these findings reveal that ferroptosis plays an important role during early infection with virulent clinical strains of Mtb by influencing bacterial spread and signaling of immune cell responders, potentially informing host-directed intervention strategies.

## Introduction

*Mycobacterium tuberculosis* (Mtb), the causative agent of tuberculosis (TB), is the leading infectious killer worldwide^1^. Over more than 10,000 years of human infection, Mtb has evolved numerous mechanisms to control the host immune response and promote disease progression^2,3^. Upon aerosol infection, Mtb first encounters and infects lung-resident alveolar macrophages (AMs)^4^. AMs remain the intracellular niche for Mtb for the first ∼10-14 days post infection (dpi), until they migrate from the alveolar space to the lung interstitium and spread Mtb to other immune responders^5^. Clinical Mtb strains that differentially regulate how and when spread from the AM occurs have been found to impact critical infection outcomes, but little is known about the mechanistic impact of these effects on downstream processes, including control of Mtb growth and downstream granuloma formation^6^. Although AM programmed cell death (PCD) is likely to facilitate spread of intracellular Mtb, how AMs initiate PCD remains poorly understood.

Numerous mechanisms of PCD have been identified, which drive distinct Mtb infection outcomes^7-9^. Broadly, these are often classified into two major categories: pathogen-restrictive apoptotic PCD and pathogen-permissive necrotic PCD pathways^9^. Necrotic PCD pathways, including necroptosis, pyroptosis, NETosis, and ferroptosis, promote the dissemination of intracellular Mtb and can elicit increased and pathological inflammatory responses^9-11^. Multiple mycobacterial effectors have been identified that promote necrotic cell death, suggesting that control over PCD is a crucial host-pathogen interface^9, 12-14^. Regulation of PCD is highly complex due to cell-lineage-specific predispositions that shift dynamically in response to the inflammatory milieu and bacterial burden^8^. Further understanding these bacterial and host effector mechanisms may reveal driving factors of infection progression.

In contrast to other innate immune responders to Mtb, AMs are characterized by a hypo-inflammatory response and early upregulation of a cell-protective transcriptional response driven by the anti-ferroptosis factor, NRF2^15-18^. Ferroptosis is a recently appreciated form of necrotic PCD in which excessive oxidative stress, iron accumulation, and the presence of polyunsaturated fatty acids (PUFAs) drives lipid peroxidation that ultimately damages cellular membranes^19,20^. In response to lipid peroxidation, NRF2 upregulates numerous oxidative stress response genes including *Slc7a11* and *Gpx4*, which regulates cysteine transport for glutathione synthesis and detoxifies lipid peroxides, respectively^19^. Loss of these genes is linked to increased necrosis and Mtb burden at later stages of Mtb infection^21^. Loss of NRF2 expression in mice increases AM PCD by 10 days post infection (dpi) with the lineage 4 H37Rv strain of Mtb, potentially highlighting AM-specific regulation of ferroptosis in early infection^18^. Possibly due to the NRF2 antioxidant response, AMs have been suggested to be relatively resistant to necrotic cell death during standard aerosol infection conditions with H37Rv^22^. However, to our knowledge, little is known about the effects of ferroptosis during the critical early stages of infection.

Recent studies have associated ferroptosis with poor bacterial control and pathological inflammatory responses during late-stage progression of pulmonary TB disease. Differences in expression of *Gpx4* and *Bach1*, a negative regulator of NRF2 activity, have been associated with human blood transcriptional signatures in individuals with active TB^21,22^. Genetic ablation of these genes in mice has been shown to significantly modulate bacterial burden as early as 17 dpi, and mouse survival after 200+ dpi. Importantly, treatment of mice with a ferroptosis inhibitor, ferrostatin-1 (Fer-1), after 14 dpi, when AMs are no longer the primary reservoir, significantly decreased bacterial burden at 28 dpi during H37Rv Mtb infection^11^. The effects of Fer-1, which acts as a potent lipid peroxide radical scavenger, suggest that modulating lipid peroxidation is an important and targetable pathway which may warrant further investigation.

Lipid peroxidation has recently been identified as an upstream inducer of type 1 interferon (IFN) signaling^23^. Early type 1 IFN signaling increases bacterial load, drives neutrophil swarming, and decreased CD4^+^ T cell infiltration^24,25^. Interestingly, these phenotypes are recapitulated during infection with the highly virulent lineage 2 Beijing strain of *M. tuberculosis*, SA161, but assessments of lipid peroxidation have not yet been performed^26-29^. The effects on lipid peroxidation and ferroptosis in host cells during infection with other clinical Mtb strains have not been described.

The above observations suggest that AM lipid peroxidation and ferroptosis during the earliest stages of infection may significantly regulate the efficacy of downstream responses. To investigate, we assessed the contribution of ferroptosis to H37Rv and SA161 infection during early timepoints when Mtb is transitioning from AMs to other innate immune cells (10-14 dpi)^5^. We show that Mtb infection induces an early pro-ferroptotic transcriptional signature in the lung, which is driven by AMs and is significantly increased by infection with SA161 relative to H37Rv. The strain-specific differences are associated with insufficient upregulation of ferroptosis suppression genes during SA161 infection, including the major lipid peroxidation suppressor, *Slc7a11. In vitro*, BMDM infection by SA161 results in significantly increased induction of lipid peroxidation and cell death in comparison to H37Rv. In C3HeB/FeJ mice, Fer-1 inhibition of lipid peroxidation for the first 14 days of *in vivo* infection led to significant decreases in bacterial burden for SA161, while H37Rv infection was unaffected. Microscopy analysis revealed that Fer-1 administration increased containment of Mtb infection within AMs, limiting its spread to monocyte-derived macrophages (MDMs). In addition, Fer-1 administration disrupted the association of neutrophilic infiltration with Mtb and their associated type 1 IFN response, which are both implicated in impairment of effective TB control^23-26^. Together, these findings add insight into the complexity of PCD during Mtb infection, demonstrating critical temporal dynamics of cell death during infection and suggesting that early induction of ferroptosis could be an important contributor of Mtb strain virulence.

## Results

### AMs have an early spike in pro-ferroptotic gene expression which is modulated by SA161

To assess cell-specific susceptibility to ferroptosis over the course of Mtb infection with different strains, we performed a transcriptional analysis of ferroptosis-related gene (FRG) expression. Transcriptomic analysis was performed on a large published single-cell RNA-sequencing (scRNAseq) dataset, which assessed lung immune cell expression with conventional dose (∼20-100 cfu) of H37Rv (17 and 34 dpi) and SA161 (10, 17, and 34 dpi) Mtb strains in both C3HeB/FeJ (C3H) and C57BL/6 (B6) mice (Fig. 1A)^30^. Prior to sequencing, enrichment for relevant immune populations was performed by flow sorting for IV-negative, and AM cells. We utilized gene set variation analysis (GSVA) to independently score relative expression of the ferroptosis driver FRG and suppressor FRG gene sets which were annotated and compiled within the FerrDB V2 database using individual pseudobulked expression per mouse^31,32^. To predict individual sample ferroptotic potential, we calculated a net FRG score by subtracting GSVA-assessed FRG-driver minus FRG-suppressor score. A high FRG score would denote high ferroptotic driver gene expression relative to compensatory ferroptotic-suppressor expression. In C3H mice, we found that net FRG score was upregulated during early infection relative to uninfected day 0 samples (10-17 dpi) (Fig. 1B). Interestingly, this increase was more prominent in SA161 infected mice, when it was significantly increased as compared to H37Rv at 17 dpi. Due to differences in cell proportions during infection progression^30^, we hypothesized that the differential net FRG score may be partially driven by the presence of different immune populations with distinct expression profiles.

**Figure 1.**
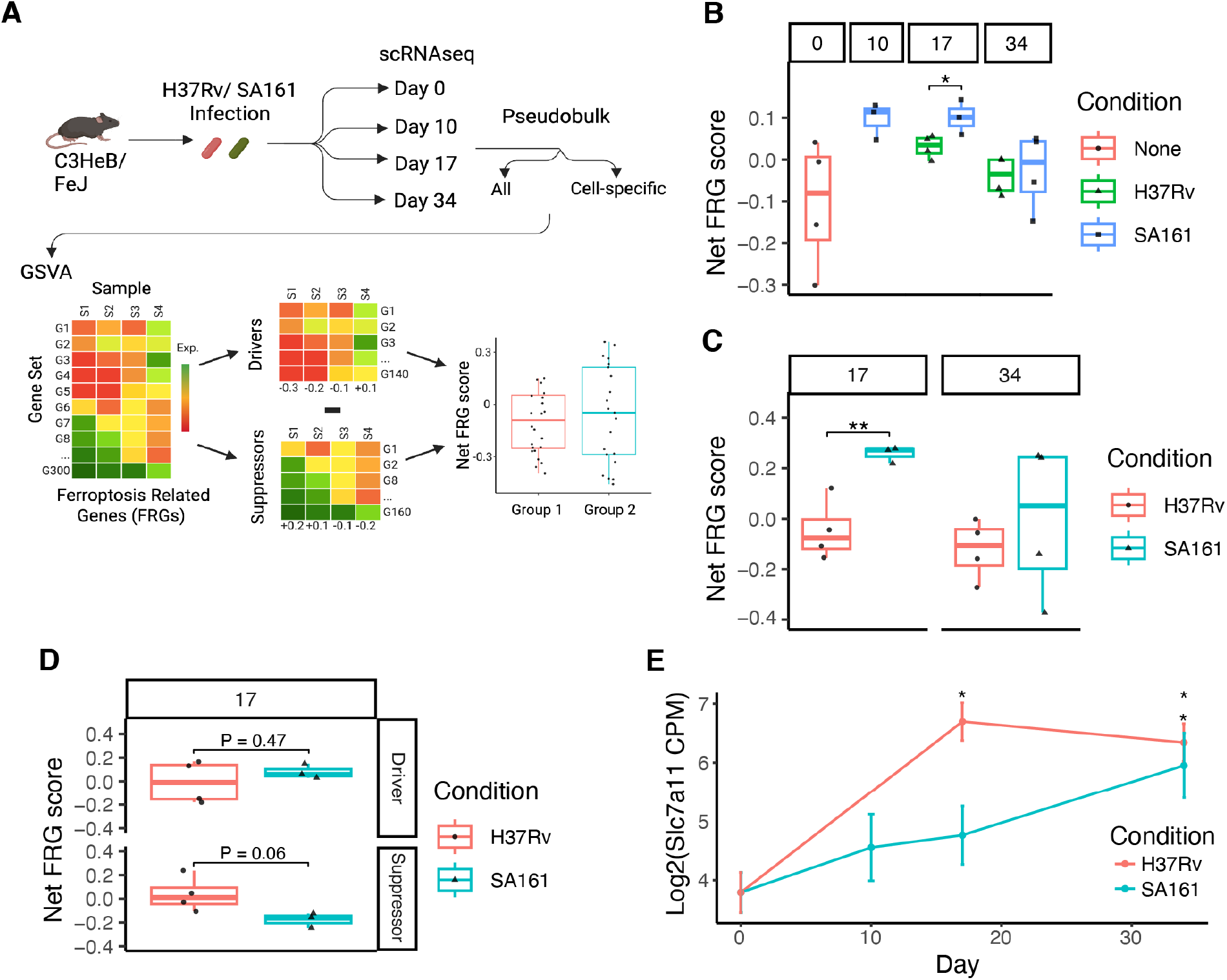
SA161 induces increased pro-ferroptotic gene expression by 17 days post infection in C3H mice. A) Diagram of GSVA analysis workflow and composition of scRNAseq dataset analyzed. Created with BioRender(Biorender.com) B) GSVA analysis identifies a significant increase in net FRG score during early infection timepoints (0-17 dpi) and is significantly higher in SA161 than H37Rv at 17 dpi. C) GSVA of AM-specific pseudobulked expression profiles reveals AM-specific increases in SA161 net FRG score over H37Rv at 17 dpi. D) GSVA of individual FRG-Driver and FRG-Suppressor scores reveals that lack of suppressor upregulation is the primary driver of net FRG scores at 17 dpi. E) Significant Mtb-induced upregulation of Slc7a11 expression is not observed for SA161 until the later 34 dpi timepoint in comparison to 17 dpi for H37Rv. Results shown are from individual pseudobulked expression of 3-4 C3H mice per timepoint and condition. Single group strain net FRG score comparisons were assessed using an unpaired t-test and timepoint comparisons were assessed using a linear model. Expression was profiled by deseq2^45^ individually at each timepoint in comparison to day 0 uninfected values after which FDR correction was performed. (*, P < 0.05; **, P < 0.01).

To assess if strain differences in net FRG score were present in individual cell populations, we re-performed GSVA on AMs which are the dominant infected cell type at day 10, when the initial increase in net FRG score was observed. Interestingly, when limited to AM expression, we observed an even greater increase in SA161-induced net FRG score in comparison to H37Rv (Fig. 1C). Interestingly, no significant FRG score differences were seen between strains in monocyte-derived macrophages, which may suggest that some early differences in strain-induced FRG transcriptional profiles are AM-driven (Supp Fig. 1A). The 17-dpi net FRG score differences were largely driven by a lack of FRG-suppressor gene set expression within SA161-infected samples (Fig 1D). Within this gene set, the most significantly differentially regulated gene was the cysteine importer *Slc7a11*, which protects against oxidative stress by supporting glutathione biosynthesis. When we assessed *Slc7a11* expression over the course of infection, we found that although *Slc7a11* was significantly induced by H37Rv at 17 dpi, *Slc7a11* induction was substantially lower during SA161 infection and not significantly increased until 34 dpi (Fig 1E). Importantly, analysis of B6 mice from the same dataset also showed a consistent increase in 17 dpi net FRG score and decreased *Slc7a11* expression during SA161 infection (Supp Fig 1B, Supp Table 1). Together, these initial analyses indicate that an increased pro-ferroptotic burden may exist during early *in vivo* infection with the SA161 strain of Mtb.

### SA161 infection induces increased levels of lipid peroxidation and cell death in BMDMs

To investigate whether the Mtb strain-dependent effects on transcriptional signatures translated to functional differences, we assessed how SA161 and H37Rv modulate ferroptosis induction kinetics during *in vitro* infection of BMDMs. We assessed for lipid peroxidation and cell death using a paired microscopy assessment of BODIPY-C11 581/591 lipid peroxidation and SYTOX dead cell staining. Upon peroxidation, the fluorescence ratio of BODIPY-C11 581/591 shifts allowing for a ratio of peroxidized BODIPY to be determined. We subjected C3H BMDMs to SA161, H37Rv, or mock infection at an MOI of 2 followed by imaging at 4, 24, 48, and 72 hours post infection (hpi). No baseline differences in lipid peroxidation between H37Rv and SA161 were observed at 4 hpi (Supp Fig. 2A). However, by 48 hpi, SA161 induced significantly more lipid peroxidation than H37Rv and this was maintained at 72 hpi (Fig. 2A, Supp Fig. 2A).

**Figure 2.**
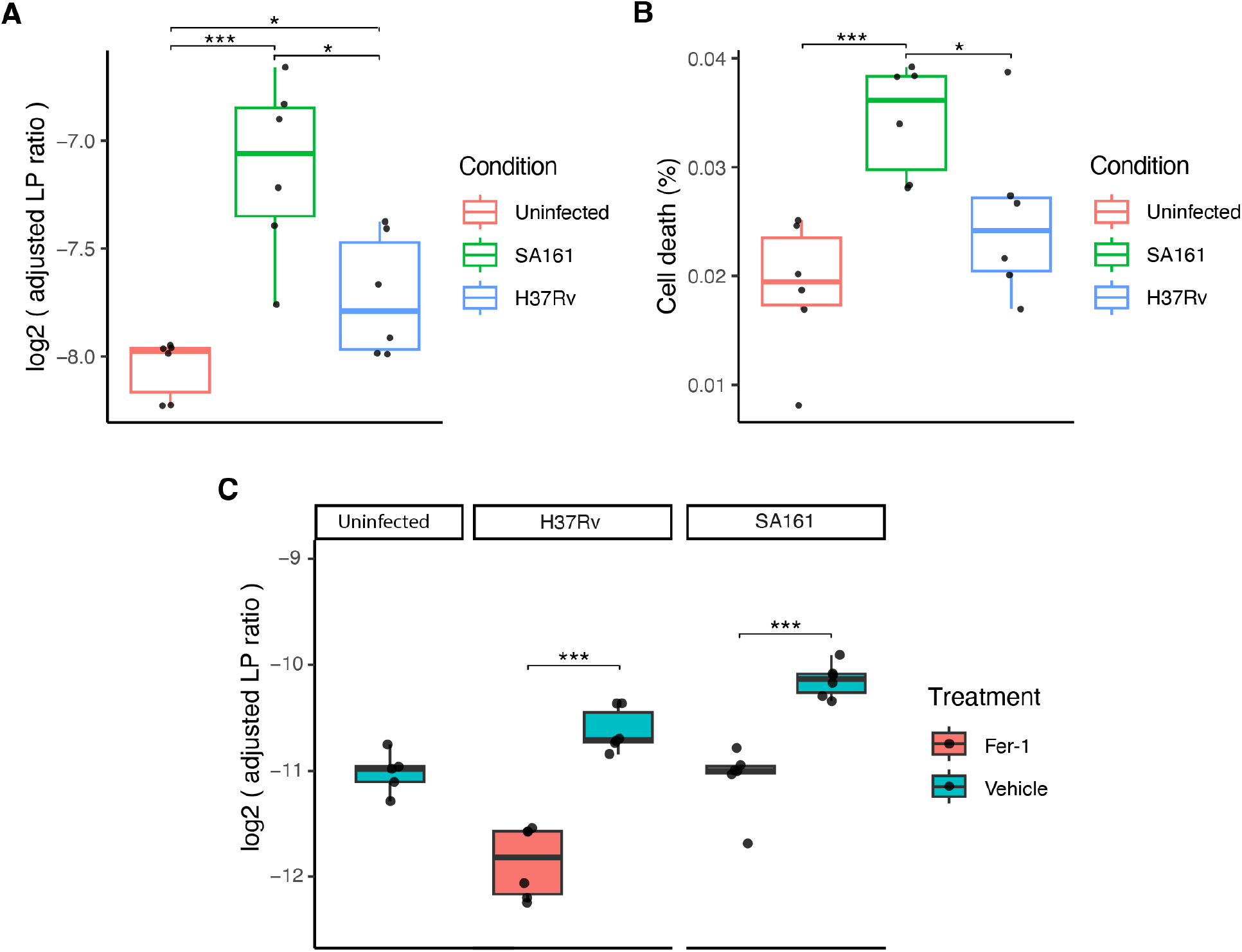
SA161 induces increased lipid peroxidation and cell death in comparison to H37Rv *in vitro* which can be modulated with Fer-1. C3H BMDMs were subjected to SA161, H37Rv, or mock infection at an MOI of 2 for 48 hours followed by bodipy-c11 581/591 or Sytox deep red staining was performed. A) SA161 infection induced an increased ratio of peroxidized Bodipy 581/591 signal in comparison to uninfected and H37Rv BMDMs indicating increased lipid peroxidation. B) Increased sytox live cell impermeable nuclear staining was observed for SA161-infected BMDMs indicating increased cell death coinciding with lipid peroxidation. C) Decreased bodipy-c11 581/591 peroxidation ratio in H37Rv and SA161 infected BMDMs upon fer-1 treatment indicates fer-1 knock down of lipid peroxidation *in vitro*.Significance was assessed using linear mixed model adjusting for the effect of order of imaging on stain development followed by unpaired t-test comparisons for individual groups. Individual data points and adjusted values are reported in Supp Table 2. (*, P < 0.05; *** P < 0.001). Results shown are representative of 6 biological replicates and are representative of A,C) 2 or B) 1 independent experiments performed.

Furthermore, the increase in SA161-induced lipid peroxidation was maintained in BMDMs from B6 mice (Supp Fig. 2B). The SA161-induced increase in lipid peroxidation at 48 hours was accompanied by a corresponding increase in cell death compared to H37Rv-infected and uninfected BMDMs (Fig. 2B). In contrast, despite a significant increase in H37Rv-induced lipid peroxidation at 48 hpi, this was not reflected in a significant increase in cell death. To assess if Mtb-induced lipid peroxidation could be modulated, we treated BMDMs with the ferroptosis inhibitor, Fer-1, starting at 4 hpi. By 48 hpi, Fer-1 treatment resulted in significant reductions in both H37Rv and SA161-induced lipid peroxidation (Fig. 2C). Overall, these findings indicate that SA161 induces increased lipid peroxidation relative to H37Rv and raises the hypothesis that this elevated lipid peroxidation could result in the observed *in vivo* transcriptional FRG signatures.

### Control of lipid peroxidation is important during early infection for SA161

Since SA161 infection increased lipid peroxidation and cell death and since this effect could be modulated by Fer-1 treatment, we next investigated the effect of Fer-1 treatment during *in vivo* SA161 and H37Rv infection (Fig. 3A). Since our transcriptomic signatures suggested early effects within AMs, we treated mice with Fer-1 (3mg/kg) or vehicle intraperitoneally for the first 13 days following conventional dose (20-100 cfu) aerosol Mtb infection. At 10, 14, and 28 dpi, mice were sacrificed and assessed for bacterial burdens, and lungs were embedded for imaging. To assess strain-induced lipid peroxidation and efficacy of Fer-1 knockdown, we performed a TBARS assay quantifying malondialdehyde (MDA) as a marker of lipid peroxidation within whole lung lysates at 10 and 14 dpi (Fig. 3B). From 10 to 14 dpi, MDA levels were highly significantly increased by H37Rv or SA161 infection, and MDA levels were significantly decreased by Fer-1 treatment (Fig. 3B). Although MDA increased from 10 to 14 dpi for both strains, at 10 dpi, SA161 infection induced significantly more MDA than H37Rv for vehicle treated mice (Fig. 3AB. To assess the impact of these lipid peroxidation differences, we examined differences in lung bacterial burdens. At 10 dpi, Fer-1 treatment within SA161 infected mice resulted in a modest but significant increase in bacterial burden compared to vehicle treated controls (Fig. 3C). Surprisingly, at 14 dpi with SA161, the effect of Fer-1 treatment reversed, and a highly significant decrease in bacterial burdens was observed (Fig. 3D). In contrast, within H37Rv infected mice, no effect of Fer-1 treatment on bacterial burdens was observed. Of note, at 14 dpi, there was no significant difference in bacterial burden between SA161 and H37Rv infected mice with Fer-1 treatment (Fig. 3D). These findings indicate that during early Mtb infection, blockade of lipid peroxidation significantly modulates bacterial burdens, specifically during SA161 infection: eliciting a detrimental effect at day 10 and a beneficial effect at day 14.

**Figure 3.**
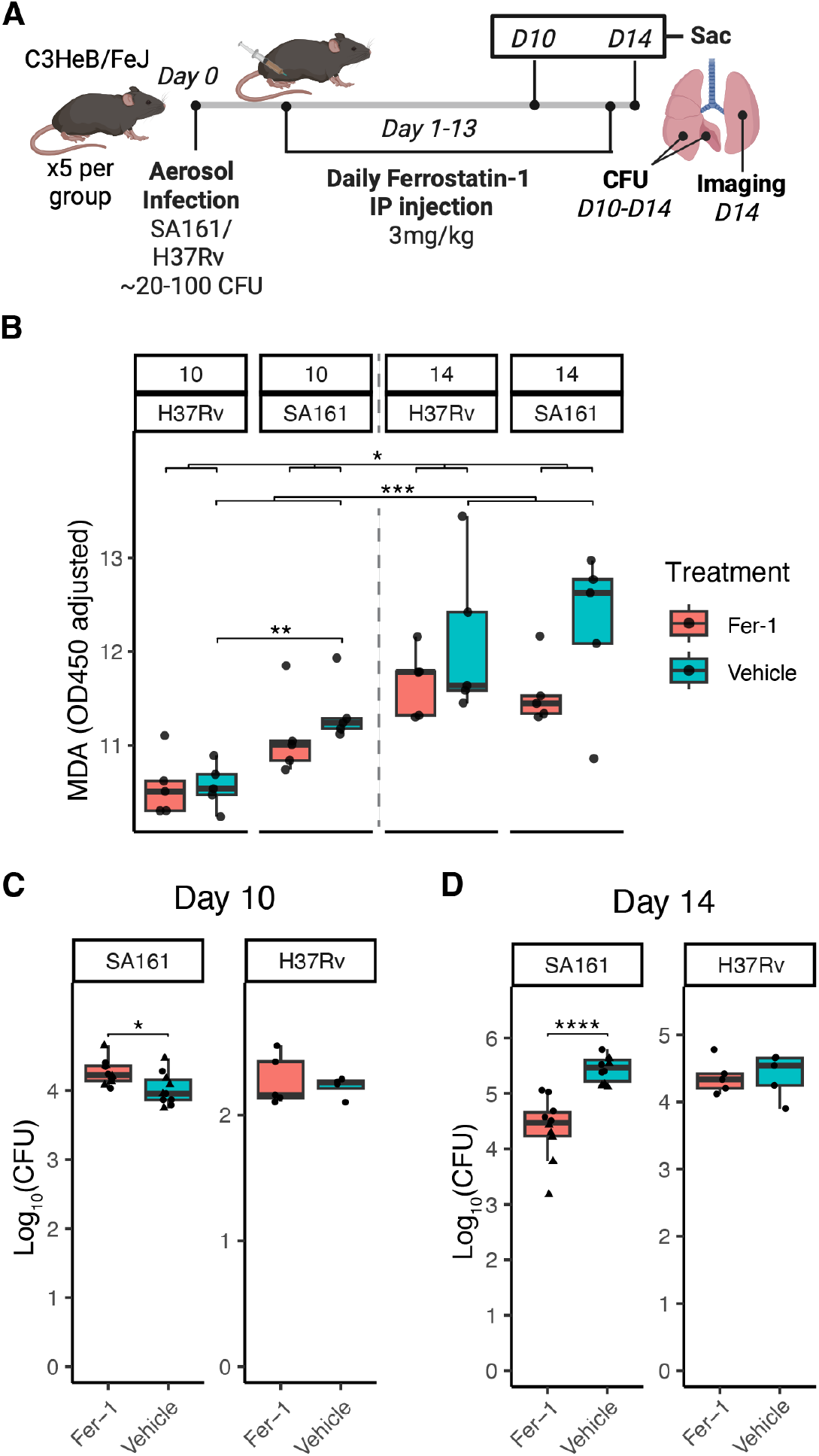
Fer-1 administration *in vivo* modulates Mtb-induced lipid peroxidation and causes SA161 specific differences in bacterial burden at 10 and 14 dpi. A) C3H mice were subjected to aerosol infection with SA161 and H37Rv followed by daily administration of Fer-1. Created with BioRender(Biorender.com). B) 10 and 14 dpi lung homogenates were assessed for MDA as a marker of lipid peroxidation revealing significant differences in lipid peroxidation from 10-14 dpi, between untreated strains at 10 dpi, and between Fer-1 treatment groups. C) Day 10 bacterial burdens in mouse lungs were assessed by CFU and an SA161-specific increase in bacterial burden upon Fer-1 treatment. D) Fer-1 results in a highly significant decrease in 14 dpi lung bacterial burdens for SA161 while H37Rv is unaffected. MDA values were adjusted using a linear model to adjust for non-specific OD450 absorbance. Individual data points and adjusted values are reported in Supp Table 2. Individual and cross group comparisons were performed using a linear model with strain and timepoint as covariates when necessary. (*, P < 0.05; *** P < 0.001, **** P < 0.0001). Results are representative of 4-5 mice per group and was independently repeated twice for SA161 or once for H37Rv and MDA assessments.

### Fer-1 inhibition decreases spread of infection from AMs and altered type 1 IFN dynamics

To investigate the potential mechanisms underlying why Fer-1 administration reduced SA161 bacterial burden by 14 dpi, we assessed infected whole mouse lung sections for pre-lesions by confocal microscopy. Infected cells were assessed by colocalization of Mtb and composition of the pre-lesion microenvironment was assessed by relative cell type proportions as well as a marker of type 1 IFN signaling (ISG15). Within Fer-1 treated mice, Mtb was often found to be restricted to AMs which were often more central to the lesion, whereas in vehicle treated mice, MDM and neutrophilic infiltration was more prominently associated with Mtb signal. (Fig. 4A). Within macrophage subtypes, Fer-1 treatment resulted in a significantly higher number of infected AMs in comparison to infected MDMs (Fig. 4B). Pre-lesion microenvironments were defined by the presence of mCherry signal, and AM, MDM, neutrophil, and Mtb abundance was quantified. Infected cells were determined by colocalization of Mtb signal within a cell. Within SA161-infected, vehicle treated mice, Mtb signal across pre-lesions was significantly correlated with neutrophil abundance (Fig. 4C). However, in Fer-1 treated samples, this neutrophil correlation was lost, and instead was significantly correlated with AM abundance. Furthermore, within the pre-lesion microenvironment, Fer-1 treatment disrupted the correlation of neutrophil and ISG15 staining which was present in control samples. In line with prior observations of bacterial burden in 14 dpi H37Rv lungs, fewer differences in pre-lesion composition were observed according to Fer-1 treatment (Fig. 4D). No significant differences in infected AM to MDM proportion were observed. In contrast to SA161 control samples, H37Rv control samples still maintained a significant correlation of Mtb with AMs across pre-lesion microenvironments (Fig. 4E). Together, these data suggest that decreased SA161 bacterial burden after Fer-1 administration correlates with increased presence of AMs which contain the Mtb infection and decreasing spread of infection to other innate responders.

**Figure 4.**
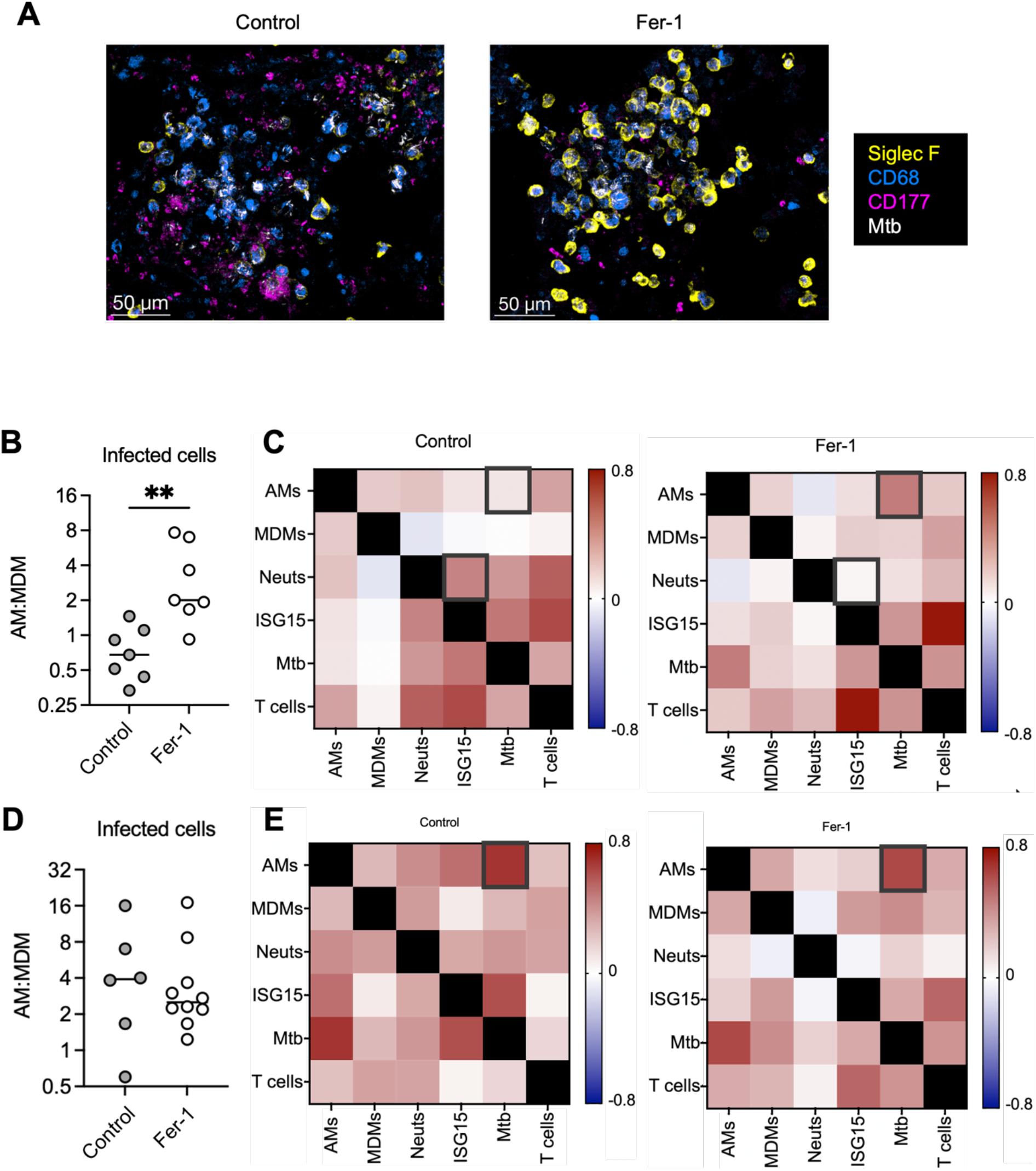
Fer-1 treatment results in containment of SA161 to AMs in 14 dpi mouse lung pre-lesions. Pre-lesions from 14 day SA161 infected mouse lungs were subjected to multi-parameter confocal microscopy. A) A representative image of pre-lesions shows increased association of Mtb labeling to AMs after Fer-1 treatment in comparison to MDMs and neutrophils in control populations. B) Ratio of infected AMs to MDMs across mouse lung pre-lesions indicates increased containment of SA161 to AMs after Fer-1 treatment. C) Composition of cell type staining was quantified across mouse lung pre-lesions indicating a shift of Mtb signal correlation with neutrophils to AMs after Fer-1 treatment and decreased correlation of neutrophils to ISG15. D) Similar correlation analyses identified that Mtb to AM correlation was maintained in Control samples within H37Rv samples. E) No significant differences in the ratio of infected AMs to MDMs was identified in H37Rv infection. Correlation assessment was performed by spearman correlation. Significance was determined by unpaired T-test. (**, P < 0.01). Images and quantification are result of multiple pre-lesions across mice from 2 (SA161) or 1 (H37Rv) independent experiments.

## Discussion

During the course of infection, Mtb establishes intracellular infections in a number of different immune cells, starting with AMs before spreading to neutrophils, monocyte-derived macrophages, monocytes, and dendritic cells^4,5^. Mycobacterial influence of the timing and mechanism of PCD to enable cell-to-cell spread within each of these cell types is essential to determining downstream infection outcomes^9-12^. Here, we have shown that the hypervirulent Lineage 2 Beijing Mtb strain, SA161, induces pro-ferroptotic transcriptional responses and excess lipid peroxidation in macrophages during early infection, relative to H37Rv. Moreover, SA161 bacterial burden at day 10 is significantly enhanced by early chemical inhibition of lipid peroxidation but also significantly attenuated by day 14 when the same pathway is blocked. This suggests that lipid peroxidation may be a virulence strategy for SA161, where the effects differ based on timing and cell-type.

During later stages of infection: 28 dpi and on, necrotic cell death has been shown to induce hyperinflammation, decrease immune responses, and reduce host survival^10,11^. At these timepoints, reducing lipid peroxidation has been shown to be effective at controlling Mtb-induced necrotic cell death^11,21,22^. However, the regulatory mechanisms and phenotypic consequences of lipid peroxidation and ferroptosis have not been explored during the timepoints when Mtb must spread from their initial AM intracellular reservoir to further propagate the infection. For example, it is not clear whether early spread out of AMs would benefit the host by accelerating the immune response or benefit the bacteria by providing additional cellular niches for replication. How each form of PCD would affect the microenvironment at this stage of infection remains undefined. Lipid peroxidation has recently been identified as an upstream regulator of type 1 IFN signaling, which is associated with neutrophilic recruitment and decreased adaptive response^23^.

Multiple studies have associated human FRG transcriptional signatures with TB disease progression during chronic disease^22,33,34^. In particular, decreased *GPX4* and increased *BACH1* expression has been identified in active TB patients in comparison to controls^21,22^. Broader assessment of ferroptotic predisposition using a GSVA-based FRG scoring method has been utilized in multiple disease contexts, including liver disease, cancer, and in TB where FRG scores from individuals with active TB were significantly different than healthy controls^33-36^. To assess ferroptosis throughout Mtb infection, we utilized a broad FRG analysis encompassing all annotated ferroptosis drivers and suppressors in the FerrDB V2 database^32^. By using a subtracted GSVA approach, our analysis allowed us to assess samples for increased ferroptotic propensity either due to elevated driver gene expression or inadequately compensatory suppressor gene response. We found that net FRG scores spiked during early Mtb infection and that this spike was significantly higher in SA161-infected mice than H37Rv by 17 dpi. Isolation of expression to AM-annotated cells further increased this stratification. Investigation into the sources responsible for the FRG score spike revealed a lack of suppression signal in genes including *Slc7a11*. H37Rv infection induced a highly significant increase in *Slc7a11* expression by 17 dpi. This is in line with prior studies of AMs which have shown that H37Rv infection induces a NRF2-driven increase in *Slc7a11* expression^15,17^. In contrast, SA161 infection did not upregulate *Slc7a11* expression until 34 dpi. Although the mechanisms of *Slc7a11* expression were not identified, it is intriguing to posit that differences might be due to SA161, lineage 2 (L2) features, in comparison to H37Rv which is lineage 4 (L4). L2 strains have previously been identified to induce significantly more type 1 IFN through increased STING activation, which is a known negative regulator of *Slc7a11* expression^37,38^. Further mechanistic assessment is necessary to determine relevance in the broader field of Mtb response biology.

When we assessed if our transcriptomic signatures validated *in vitro*, we found that SA161 significantly increased lipid peroxidation and cell death over H37Rv by 48 hours of infection. Our results with low MOI H37Rv infection align with previous BMDM studies in which H37Rv infection at an MOI of 1 resulted in no significant increase in cell death over 4 days and induced only a small increase in lipid peroxidation^11^. Our *in vitro* data suggest that, unlike H37Rv, SA161 may have intrinsic ferroptosis-inducing properties in physiologically relevant MOI. Although, a recent study suggested mouse strain-specific differences in lipid peroxidation from C3H and B6 BMDMs stimulated with TNF^23^, our lipid peroxidation findings were consistent across mouse strains. Even though our transcriptomic signature was found to be specific to AMs, we also detected elevated lipid peroxidation upon infection in BMDMs *in vitro*. Further assessment of mouse and bacterial strain-dependent effects on lipid peroxidation and ferroptosis in AMs is warranted.

Mimicking our *in vitro* studies, both H37Rv- and SA161-infected mouse lungs increased in lipid peroxidation over time and was significantly suppressed with Fer-1 treatment. However, SA161 induced significantly more lipid peroxidation by 10 dpi and effects of Fer-1 knockdown on bacterial burden were specific to SA161. At 10 dpi, we found that Fer-1 inhibition increased SA161 bacterial burden. Interestingly, this may align with prior work in which control B6 mice had decreased AM cell death and higher bacterial burden in comparison to those with AM-targeted NRF2 deletion^15^. Since several studies have identified day 10 as a timepoint just before AMs begin to migrate to the lung interstitium, our data supports a model in which AM cell death is only beneficial prior to this migration^5^. Due to AMs being a relatively permissive niche for Mtb, we hypothesize that it may benefit Mtb to grow withinAMs before they traverse to the interstitium, where cell death can drive greater recruitment and spread to other responding phagocytes^15-18^. In line with this hypothesis, AM depletion prior to infection has previously been shown to increase ability to control bacterial burden^39^.

In contrast to 10 dpi, Fer-1 inhibition of lipid peroxidation resulted in a substantial decrease in bacterial burden at 14 dpi with SA161. This decrease was specific to SA161 with no effect observed in H37Rv. This suggests that increased bacterial burden associated with SA161 hypervirulence may be mitigated by lipid peroxidation. Further investigations into the impact of inhibiting lipid peroxidation on overall SA161 virulence are warranted. Although the effect of Fer-1 treatment on bacterial burden by 14 dpi is in line with previous research of Fer-1 treatment from 14-28 dpi, the SA161 specificity and the host cell types involved at this time point suggest that different mechanisms may be involved^11^. From 10-14 dpi, AMs facilitate spread of contained intracellular bacteria to responding phagocytic neutrophils and monocyte-derived macrophages (MDMs) and transition away from being the majority infected cell type^5^. As such, AM cell death is likely an essential aspect of this transition which may be modulated by Fer-1 at this timepoint. In line with these hypotheses, our imaging suggests that Mtb within SA161-infected mouse lungs at 14 dpi are more highly contained within AMs following Fer-1 treatment. This AM containment coincides with decreased infection of MDMs and neutrophilic infiltration.

Furthermore, upon Fer-1 treatment, correlation of neutrophil and ISG15 abundance was lost. ISG15 is a type 1 IFN-inducible gene and can act as a marker of type 1 IFN signaling^40^. Type 1 IFN signaling and neutrophil abundance are associated with negative infection outcomes, driving a detrimental recruitment and NETosis loop which can increase bacterial burden and hinder adaptive response recruitment^25^. Overall, these data support our hypothesis that decreasing SA161-induced lipid peroxidation may limit spread of intracellular Mtb from within infected AMs and decrease bacterial burden and neutrophilic infiltration which are characteristic of SA161 infection.

This study focused on characterizing only two Mtb strains, and further work to contextualize our findings to the broader landscape of Mtb strain physiology is essential. Further research is necessary to determine if increased lipid peroxidation is an SA161-specific phenotype, or if it more broadly applies to other hypervirulent or lineage 2 strains. Additionally, this study largely focused on lipid peroxidation as a marker of ferroptotic cell death. Although, lipid peroxidation-associated cell death is characteristic of ferroptosis, necrotic cell death pathways often coincide and can act in concert^41^. We sought to combat this difficulty with consistent bioinformatic, *in vitro*, and *in vivo*, investigations. However, further mechanistic investigation is necessary to isolate specific PCD effects. The mechanisms by which SA161 drives increased lipid peroxidation have not yet been investigated. Although other Mtb effectors have been identified to perturb host lipid peroxidation, these act outside of the early timepoint investigated^14,42,43^. As such, SA161-induced lipid peroxidation may have unique mechanisms which will be the target of future research. Finally, the fact that Fer-1 treatment has opposing effects at day 10 and 14 suggests that ferroptotic cell death can be either beneficial or harmful to the host during Mtb infection depending on the infected cell type and the local environment, but understanding the mechanisms underlying these divergent effects will require additional studies. Further understanding of how Mtb modulates host cell PCD to control infection trajectories may yield important insight into mechanisms of strain virulence and heterogeneous host infection outcomes and inform future TB eradication efforts.

## Materials and Methods

### Mice

C3HeB/FeJ and C57BL/6 were purchased from Jackson laboratories (Bar harbor, ME). All mice were maintained in individually ventilated cages in pathogen-free conditions with a maximum of 5 mice per cage. Animals were maintained in an accredited ABSL3 facility at Seattle Children’s Research Institute with negative pressure ventilation, free access to food and water, 12hr light and dark cycles, temperature control between 22°C and 25°C, under care of full-time staff. All animals were normal health and immune status with no prior treatments or invasive studies. All studies were approved by the Seattle Children’s Research Institute Animal Care and Use Committee in adherence with the National Institutes of Health Guide for the Care and Use of Laboratory Animals. Sex and age matched female, 8-12 week old mice were used for subsequent investigations and sex was not assessed as a biological variable.

### Single cell RNA sequencing

Single cell RNA sequencing data was performed on previously published data^30^. Briefly, mouse lungs from uninfected, 10, 17, and 34 dpi mice were excised and subjected to 2x gentle homogenization using a gentleMacs dissociator (Miltenyi Biotec). To allow for removal of IV+ cells, mice were previously anesthetized and injected with anti-CD45.2 antibody intravenously. Homogenates were filtered with a 70uM strainer subjected to RBC lysis in RBC lysis buffer (Thermo), and resuspended in FACS buffer to generate a single-cell suspension. Samples were analyzed via FACS Ariall (BD) sorter to obtain autofluorescent AMs and IV-cell populations which were then recombined. Cells were processed according to 10x Genomics pipeline recommendations. Final samples were subjected to 2x 0.2uM (SpinX-CoStar) filtration prior to BSL3 removal and were submitted to Psomagen (Rockville, MD) for NovaSeq sequencing at 300M reads per sample.

### Single cell RNA sequencing processing

Single cell RNA sequencing data was processed as previously published^30^. Briefly, single-cell RNAseq sequence reads were mapped to the 10X Genomics pre-built mouse reference genome mm10-2020-A. Cells were assigned individual cell barcodes and processed with the Seurat R package for QC filtering and integration^44^. Initial filtration required cells with greater than 500 distinct genes and fewer than 5% mitochondrial genes and genes with more than 3 cells expressed. Seurat was used to correct for batch and align cells across all dataset conditions. For cell type annotation, cells were analyzed and assigned using the CellTypist python package using the CellTypist Mouse Atlas Annotations cell type model^45^. For all downstream analyses, only unvaccinated samples were assessed and all coMtb samples were removed. Pseudobulk analysis was performed using Presto, per whole sample or after CellTypist annotation specific extraction, on genes with expression in less than 100 cells to avoid effects from rarely expressed genes. Pseudobulked expression was then normalized in CPM within each sample.

### Gene Set Variation Analysis

GSVA was conducted using the GSVA package in R^31^. FRG driver and suppressor gene sets were extracted from FerrDB V2 dataset for mice^32^. If individual genes were located in both gene sets, the gene was assigned to the gene set with the highest confidence score, or removed if no decision could be made. GSVA scores were calculated across all samples of interest for each analysis independently for driver and suppressor gene sets, after which a net FRG score was calculated from driver minus suppressor score.

### Differential expression analysis

Differential expression analysis of FRG gene set expression was calculated using pseudobulk DESeq2 in R with singular comparison of log fold change gene expression for each dpi timepoint with day 0^46^. Pseudobulked expression was assessed from CellTypist assigned Alveolar Macrophages which was then counts per million and log2 normalized.

### Primary Murine Macrophage Isolation and Culture

Murine BMDMs from C3HeB/FeJ and C57BL/6 were extracted using the following protocol. Marrow from femurs and tibia was extracted via flushing with RPMI-1640 (Fisher) and single-cell suspended via pipetting. Cells suspensions were then pelleted, and resuspended in cell culture media composed of RPMI-1640 with 10% FBS (Gibco), supplemented with 2mM Glutamine, and penicillin (100 units/mL, Gibco), Streptomycin (100 ug/mL, Gibco), and 50ng/mL human MCSF (Sino Biological). Cells were plated in 100mm suspension culture dishes and allowed to differentiate for 6 days. 50% additional cell culture media was added to dishes on day 3. On day 6, non-adherent cells were removed with 3x wash with DPBS (Gibco). Remaining BMDMs were then detached with gentle pipetting following 10 minute incubation at 37c in DPBS +1mM EDTA. Cells were then resuspended in cell culture media without antibiotics, counted and plated in 96 well plates at 80,000 cells per well and allowed to rest for 1 day prior to experimentation.

### Bacterial Culture

SA161 and H37Rv Mtb infection cultures were grown from previously frozen stocks which were thawed and subcultured for 3 days and then back-diluted to an Optical Density at 600nm (OD) of 0.2 1 day prior to experiments. Strains were grown in middlebrook 7H9 Broth supplemented with 10% OADC, 0.2% glycerol, 0.1mM proprionate, and 0.05% tyloxapol to maintain extracellular Mtb factors as previously described^47^.

### In vitro macrophage infection

Infections were performed using a modified previously published protocol^48^. SA161 and H37Rv infection cultures were prepared as described above and were grown to mid-log phase (0.3-0.4 OD) and final ODs were obtained. Equivalent bacterial cultures were then pelleted at 3000xg resuspended to equivalent CFU/mL per strain. Two mLs of inoculum was then removed and subjected to 15x passaging through a 25-gauge needle for single-cell resuspension.

Resuspended cells were then diluted to in RPMI-only and added to individual wells at an MOI of 2 in order to increase likelihood of a consistent infection of at least 1 Mtb per BMDM. Infected 96 well plates were then centrifuged at 300xg for 5 minutes to facilitate Mtb descent to BMDM layer. Infection was allowed to proceed for four hours, after which infection inoculum was removed, cells were washed 3x with DPBS, and fresh cell culture media was added. To control for media usage, 50% media was removed and replaced with fresh cell culture media daily.

### *In vitro* imaging of lipid peroxidation and cell death

*In vitro* lipid peroxidation and cell death was assessed using a microscopic measurement of BMDMs following staining with Bodipy-c11 581/591 (Invitrogen) lipid peroxidation dye and Sytox deep red (Invitrogen) live cell impermeable nuclear stain. Two hours prior to imaging timepoint, lipid peroxidation positive control wells were subjected to cumene hydroperoxide (Invitrogen) at a final concentration of 100uM. Thirty minutes prior to imaging, BMDMs were then stained with a final concentration of Bodipy-c11 581/591 (10uM) and Sytox deep red (0.5uM) per well. At imaging timepoint, all cells were then washed 3x with DPBS and replaced with RPMI-only. Cells were imaged using an imageXpress micro confocal high content imaging system, with a 20x objective at 9 fields per well.

### Image processing and analysis of lipid peroxidation and cell death

All image analysis was performed using the MetaXpress image analysis software. For lipid peroxidation, total Bodipy-c11 581/591 signal was ascertained with the Texas Red channel, and cellular objects were identified using the find blobs algorithm thresholded to a width of 5-30 uM at an intensity of 1000 over local background. Oxidized Bodipy-c11 581/591 cellular objects were then identified in the FITC channel within a mask of the above objects. Vignetting within the FITC channel was adjusted for using the addition of a gaussian blur as previously described^49^. Prior to image analysis, any cellular debris was identified in the FITC channel using a find blob filter of large objects 50-500 uM in size, and these objects as well as image borders 30uM from edge were removed. Fluorescent intensity from image objects was then background adjusted to minimum local background within 50uM. Summed FITC fluorescent intensity was divided by summed Texas Red signal within each image and mean of 9 images per well was used for final lipid peroxidation ratio. For cell death analysis, total cell number was enumerated using the total Bodipy-c11 581/591 mask objects as above. Within these cells, nuclear specific staining was assessed for Sytox deep red within the Cy5 channel within this total cell mask. Sytox positive nuclei were determined by subtracting Texas Red from Cy5 image intensities showing nuclei specific staining and then counted using the find blob algorithm thresholded to a width of 3-12uM. Analysis data was then exported and processed in R. Due to the large number of images per well, and the assay being in live cells, we observed a highly significant effect of image order on cell staining intensity, likely due to continued development of wells as imaging occurred. To control for this effect, image order was adjusted for using a linear model adjustment of order as a linear effect and adjusted metrics were reported.

### Aerosol Infections

Infections with Mtb SA161 and H37Rv strains was performed, as described previousl^50^. For each infection, mice were placed in a Glas-Col aerosol infection chamber, and ∼20–100 CFUs of corresponding strain were deposited into their lungs. Infectious inoculum was then assessed from two mice per infection which were euthanized on the day of infection, and their lung homogenates were plated onto 7H11 plates for CFU enumeration.

### Ferrostatin-1 inhibitor treatment

Fer-1 knockdown *in vitro* was performed by applying Ferrostatin-1 (MedChemExpress) at a final concentration of 10uM within culture media starting after infection inoculum removal at 4 hours. To account for Fer-1 degradation, further Fer-1 was applied daily by treating with equivalent concentrations within the daily 50% media change. For *in vivo* Fer-1 treatment, Fer-1 was sequentially resuspended in DMSO, PEG-400, and PBS to a final concentration of 0.3% DMSO, 40% PEG-400, and 59.7% PBS and administered at 3mg/kg intraperitoneally daily. All Fer-1 administrations were assessed in comparison to identically performed vehicle controls with DMSO only in place of Fer-1.

### *In vivo* assessment of lipid peroxidation

Lipid peroxidation was assessed *in vivo* using the TBARS assay kit (Cayman Chemicals) for MDA content. Mouse lungs were excised and homogenized in a gentleMACS M tube (Miltenyi Biotec) containing PBS + 0.05% tween-80, placed on ice, and frozen at −80°c for subsequent analysis within 30 days. For analysis, lysates were thawed on ice, processed according to colorimetric protocol, and read at 530nM. Non-specific signal was simultaneously ascertained at 450nM. 530nM reads were highly correlated to 450nM reads due to heterogeneity of lung homogenate even more strongly than total protein content assayed using Pierce BCA assay and was likely due to differences in lysate blood content upon homogenization. To account for this sample diversity, 530nM signal was adjusted for 450nM signal using linear model regression of 450nM signal as a linear effect.

### CFU enumeration

Mouse lungs were excised and homogenized in a gentleMACS M tube (Miltenyi Biotec) containing PBS + 0.05% tween-80. Resulting homogenates were then diluted and plated onto 7H10 plates and incubated at 37°C for a minimum of 21 days prior to CFU enumeration.

### Confocal microscopy

Mouse left lungs were removed and placed in BD Cytofix (BD) and diluted 1:3 PBS for 24hr at 4°C prior to BSL3 removal. Lungs were then washed 2x in PBS and incubated in 30% sucrose for 24hrs at 4°C. Lungs were embedded in OCT and frozen using a dry ice slurry with 100% ethanol. Serial 20uM sections were taken using a Leica CM1950 cryostat. For image staining, sections were rehydrated in 0.1M Tris for 10 minutes, incubated in blocking buffer (0.1M TRIS + 1% normal mouse serum, 1% bovine serum albumin, and 0.3% Triton-X100), and then stained overnight at room temp with fluorescently conjugated antibodies. Stains were then washed with 0.1M Tris for 30 minutes and cover-slipped with Fluoromount G mounting media (Southern Biotech). Confocal images were acquired using a Leica Stellaris 8 confocal microscope.

### Histo-cytometry

Histo-cytometry analysis was performed as previously described with minor modifications^51^. Confocal images were first corrected for fluorophore spillover. Single-color controls were prepared by mixing fluorophore-conjugated antibodies with Fluoromount G mounting medium (SouthernBiotech), placing on a slide, and collecting images using the same settings as for tissue imaging. Next, the fluorophore spillover was calculated and corrected utilizing the Channel Dye Separation module in LAS X (Leica). Cell objects were created on nuclear staining using the Imaris surface object module. Object statistics were exported in CSV format. Object statistics were then imported into FlowJo software for cellular gating. Spatial analysis was subsequently performed using CytoMAP^52^. In brief, the positions of all cell objects within tissues were used for virtual raster scanning with 50-µm-radius neighborhoods. Lesions were defined by the presence of mCherry signal in a neighborhood. Cellular abundance correlations were completed using CytoMAP.

### Statistical Analysis

Statistical tests were performed in R and were selected based on appropriate assessments of distribution and variance characteristics. Exact details of tests performed are located within figure legends. Statistical differences of *in vitro* and *in vivo* assays were assessed using a two sample-t test using the lm package with adjustment for covariates when necessary^53^. When necessary, control for multiple comparisons was performed using FDR adjustment of base p-values.

## Acknowledgments

We gratefully acknowledge Alan Sher and Eduardo Amaral for helpful discussions.

## Supplemental Figures

**supplementary Figure 1.**
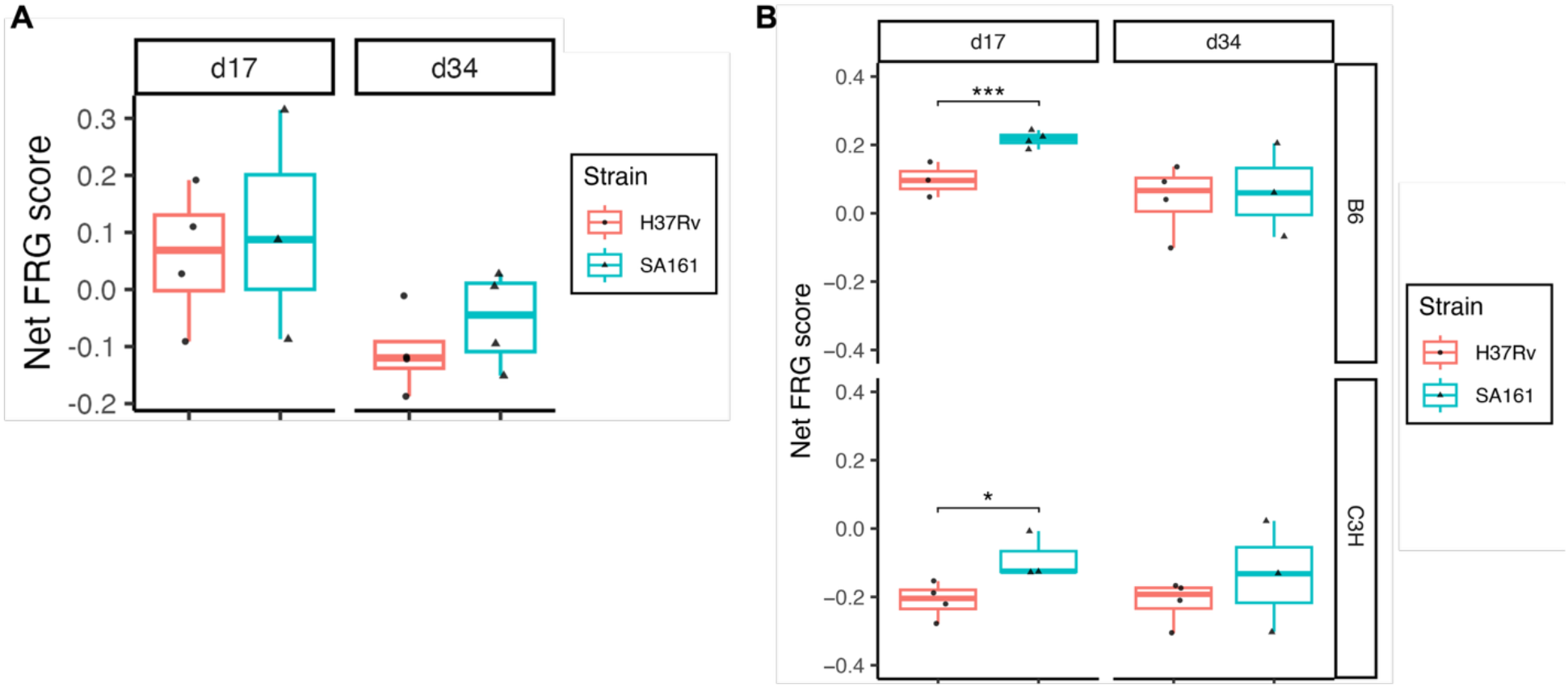
Observed 17 dpi net FRG score differences are not due to MDM expression and are consistent in B6 mice. Reanalysis of pseudobulked expression profiles. A) MDMs in C3H mice at 17 and 34 dpi show no significant differences in net FRG score. B) AM pseudobulked expression across C3H and B6 mice reveals a consistent increase in SA161 infection net FRG score at the day 17 timepoint. Single group strain net FRG score comparisons were assessed using an unpaired t-test and timepoint differences were assessed with a linear model adjusting for strain as a covariate. (*, P < 0.05).

**supplementary Figure 2.**
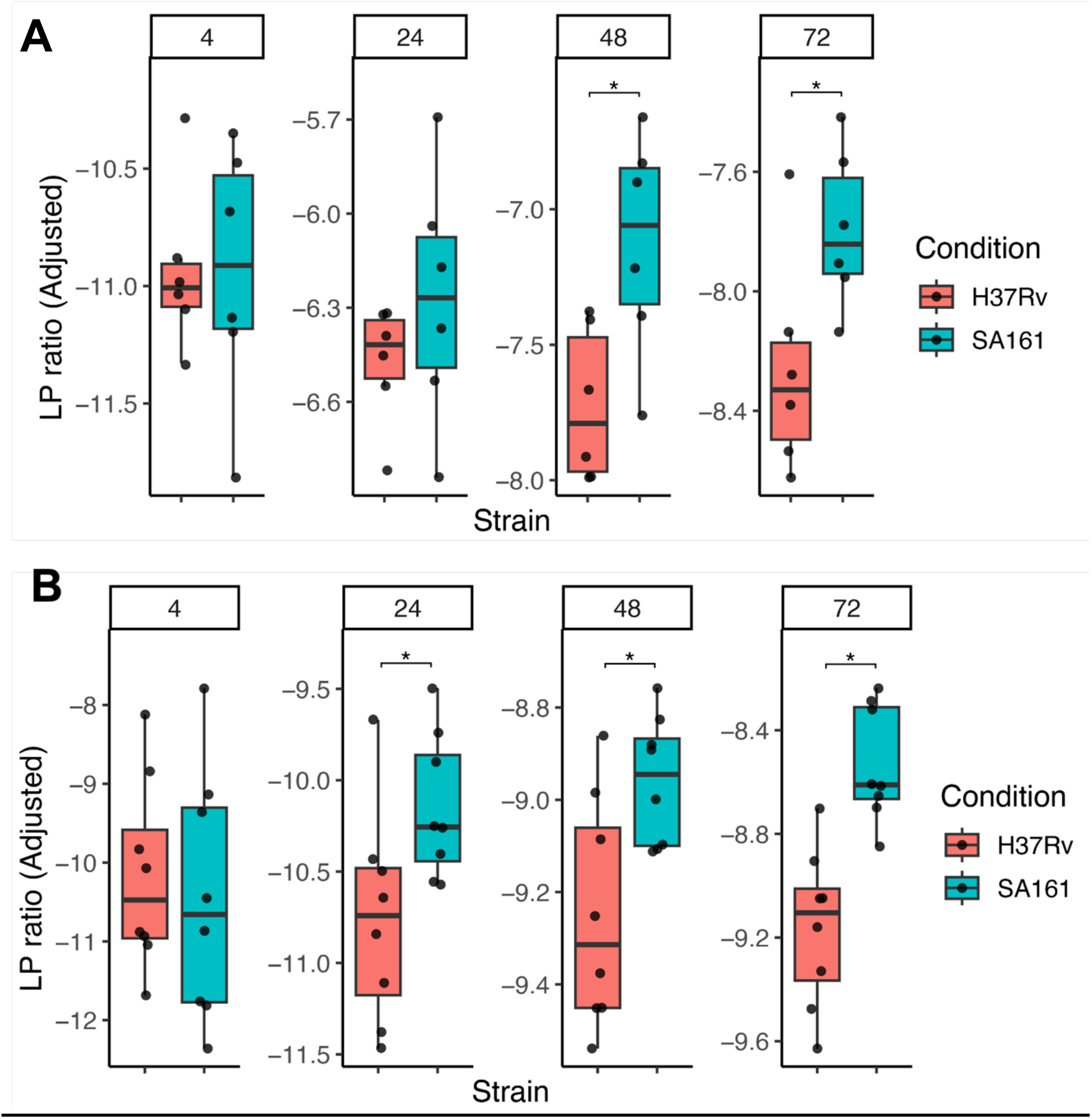
SA161 induces increased lipid peroxidation over 72 hours in C3H and B6 mice. A) C3H and B) B6 BMDMs were subjected to SA161 and H37Rv infection at an MOI of 2 for profiled daily for 72 hours following bodipy-c11 581/591 staining was performed. Significant differences according to strain were observed by 48 hours for both mouse strains and was maintained significantly by 72 hours. No differences were observed baseline at 4 hours. Significance was assessed using linear mixed model adjusting for the effect of order of imaging on stain development followed by unpaired t-test comparisons for individual groups. Individual data points and adjusted values are reported in Supp Table 2. (*, P < 0.05; *** P < 0.001). Results shown are representative of 6 biological replicates per condition.

## Supplemental tables

**Supplemental table 1) SA161 and H37Rv show distinct patterns of Mtb-induced expression over time**. Log fold change and FDR corrected P-value shown for each strain and infection time point relative to day 0 uninfected samples.

**Supplemental table 2) Plotting values with covariate adjustment**. All plotted values as unadjusted and, if necessary, their covariate assessed adjustment and adjusted values.

